# Inhibition of lysine deacetylase activity impacts formation of the vitamin D receptor activation complex

**DOI:** 10.1101/2025.02.03.636272

**Authors:** Shannon Dwyer, Kevin Kemper, Vincent J. Manna

## Abstract

The vitamin D endocrine system is responsible for the regulation of many biological processes including bone metabolism, calcium homeostasis, cell proliferation and cell differentiation. Alterations to the vitamin D signaling pathway are associated with several diseases including bone diseases, diabetes, cardiovascular diseases, autoimmune diseases, and cancer. Vitamin D precursors are obtained through diet or synthesized in the skin and must be further chemically modified to become the biologically active hormone, calcitriol. Calcitriol binds to the vitamin D receptor (VDR), a member of the nuclear hormone receptor (NHR) superfamily. VDR forms a heterodimer with retinoid X receptor (RXR) and together they bind to promoters containing vitamin D response elements (VDREs) to activate transcription of target genes. Other NHRs have been shown to accept post-translational modifications that can either increase or decrease their transcriptional output through alterations in protein-protein or protein-DNA interactions. We have generated evidence that two lysines on VDR may be targets of post-translational modifications, and alterations to lysine deacetylase activity will impact VDR transcriptional output through changes in co-activator and co-repressor binding. Together, these data suggest a novel way for the cell to modulate the response of VDR to available vitamin D.

## INTRODUCTION

Traditionally, vitamin D signaling is most known for its role in skeletal integrity, especially regulating bone metabolism as well as calcium and phosphorus homeostasis (Hausler et al., 1998). The active vitamin D hormone is responsible for intestinal absorption and renal reabsorption of both calcium and phosphorus, as well as the formation of osteoblasts, a major component vital for bone formation and remodeling (Nakamichi et al., 2017). However, the role of vitamin D extends much further than just bone health as its receptor is expressed in nearly every cell type in the body. It is involved in a wide variety of biological processes including cardiovascular health, regulation of immune response, cell proliferation and cell differentiation (Nagpal et al., 2005). Alterations to the vitamin D signaling pathway are associated with several diseases including bone diseases, diabetes, cardiovascular diseases, autoimmune diseases, and cancer. Specifically in cancer cells, vitamin D has been shown to have anti-proliferative effects via G0/G1 cell cycle arrest, induction of apoptosis, or inhibition of angiogenesis (Fujioka et al., 1998).

Active vitamin D3 (Calcitriol) is synthesized through a series of reactions to become the biologically active hormone. Pre-vitamin D3 precursors can be obtained through diet or synthesized in the skin from 7-dehydrocholesterol (Pro-vitamin D) following exposure to UVB sunlight. Pre-vitamin D3 undergoes isomerization to form vitamin D3 (Cholecalciferol). Vitamin D3 is then transported to the liver via vitamin D-binding protein (DPB) where it is hydroxylated by enzyme 25-hydroxylase (CYP27A) to form 25-hydroxycholecalciferol [25(OH)D3] (Calcidiol/pre-active vitamin D). In the kidney, the enzyme 1α-hydroxylase (CYP27B1) catalyzes the second hydroxylation to form the active hormone, 1,25(OH)2D3 (Calcitriol/active vitamin D/aD3) (DeLuca, 2004). Conversely, CYP24A1 catalyzes the degradation of both pre-active vitamin D and aD3 and is transcriptionally activated by aD3, thus regulating its levels in the body via a negative feedback loop (Jones et al., 2012). Moving forward, active vitamin D/Calcitriol, will be referred to as “aD3” for simplicity.

aD3 functions through its protein receptor, the vitamin D receptor (VDR). VDR is a member of the nuclear hormone receptor (NHR) superfamily, a group of ligand-inducible transcription factors that function as heterodimers and interact directly with DNA response elements on target genes. NHRs are comprised of an N-terminal transcriptional activation function (AF-1), a conserved DNA-binding domain (DBD), a hinge region, and a ligand binding domain (LBD). The AF-1 domain is highly variable across NHRs and is responsible for ligand independent function (Aranda & Pascual, 2001). The cysteine rich DBD consists of two zinc fingers and two α-helices that are crucial for recognition and binding to the DNA response elements. The first α-helix is responsible for binding to specific DNA bases in the major groove. The hinge region serves as a linker between the DBD and LBD and facilitates orientation of the DBD. The LBD is made up of twelve α-helices (H1-H12) and is important for binding to the NHR’s respective ligand as well as facilitating heterodimerization with partner NHRs. The LBD contains an activation function domain (AF-2) within H12 that directly interacts with the ligand. VDR’s heterodimeric partner is Retinoid X Receptor (RXR), which also serves as the binding partner for many of the other NHRs (Wurtz, 1996). In the absence of aD3, VDR forms a heterodimer with RXR and various co-repressors, such as SMRT, Nuclear receptor co-repressor (N-CoR), and Alien (Baniahmad, 2005; Dressel et al., 1999). When aD3 binds to VDR, it triggers a conformational change forming a protein interaction surface on H12 of the AF-2 domain to facilitate biding of co-activators such as SRC1, SMAD3, DRIP/TRAP complex, and CBP/p300 (Kamei et al., 1996; Rachez et al., 1998; Tagami et al., 1998; Yanagisawa et al., 1999). This complex moves to the nucleus where it transcribes genes whose promoters contain vitamin D response elements (VDREs). VDREs typically consist of DR3 motifs, which are direct repeats of a hexamer motif (AGGTCA, GGTCCA, or GGGTGA) separated by three nucleotides (Umesono et al., 1991). VDR activity modulates thousands of genes in the human genome, inducing a wide array of biological effects.

Protein post-translational modifications such as phosphorylation, methylation, acetylation, SUMOylation, and ubiquitination modulate a wide variety of processes in the cell, including protein signaling, affinities for interacting partners, localization, and degradation (Shvedunova & Akhtar, 2022). For decades, lysine acetylation was studied exclusively on histone proteins until the 1980s and 1990s when it was discovered that non-histone proteins such as α-tubulin and p53 can be targets of post-translational lysine-acetylation (Gu & Roeder, 1997; L’Hernault & Rosenbaum, 1985). Since then, thousands of acetylated lysine residues on non-histone proteins have been revealed via proteome-wide studies (Kori et al., 2017). In this reversible reaction, an acetyl group is transferred from acetyl co-enzyme A (acetyl-CoA) onto the ε-amino group of a lysine, catalyzed by lysine acetyltransferases (KATs). Conversely, this acetyl group can be removed by lysine deacetylases (KDACs) (Drazic et al., 2016). KATs are broken up into three families: GCN5-related N-acetyltransferases (GNAT) (GCN5 and PCAF), p300/CREB-binding protein (p300 and CBP), and MYST (MOZ, Ybf2, Sas2 and Tip60). KDACs are separated into four classes: class I (KDACs 1, 2, 3, 8), class II (KACs 4-7, 9, 10), class III (Sirtuins 1-7) and class IV (KDAC11). The enzymatic activity of classes I, II, and IV are Zn2+-dependent whereas the class III Sirtuins are nicotinamide adenine dinucleotide (NAD+) -dependent (Ali et al., 2018). Lysine acetylation plays a role in modulating protein function through a variety of ways. The addition of the acetyl group onto the lysine neutralizes the residue’s positive charge, therefore affecting its electrostatic interactions with other binding partners. In addition, the modification may also alter the protein’s conformation, stability, localization, or ability to accept other post-translational modifications (Shvedunova & Akhtar, 2022). Eleven different NHRs have been shown to be directly acetylated (Wang et al., 2011). Some of these acetylation sites act as positive regulators of transcriptional activity, whereas other acetylation sites act as negative regulators. For example, both the Estrogen receptor (ER) and the Androgen receptor (AR) undergo acetylation at lysine residues within the hinge region that increase ligand-induced transcriptional output (Wang et al., 2001). Alternatively, when the Farnesoid X receptor (FXR) gets acetylated at two lysine residues in the DBD and hinge region, FXR and its heterodimeric partner, RXR, are unable to bind to DNA and transcriptional activity decreases (Kemper et al., 2009). Studies have shown that altering the transcriptional output of NHRs outside of their normal range, whether positively or negatively, can contribute to the progression of various diseases and cancers (Anbalagan et al., 2012).

Investigating potential VDR acetylation sites and the impact of acetyltransferase activity on VDR transcriptional activity may reveal novel mechanisms of VDR transcriptional regulation and may provide insight into the pathogenesis of some of these diseases and cancers.

## Materials and Methods

### Plasmids

The VDR reporter plasmid, CYP24A1-FLASH, was generated by modifying an existing beta-catenin reporter, TOP-FLASH (Addgene plasmid #12456). The CYP24A1 promoter was cloned from human nasal epithelial cells (HNECs) from position -1500 to -25 regarding the transcription start site. The CYP24A1 promoter was inserted into the TOP-FLASH reporter upstream of the firefly luciferase coding sequence using Kpn1 and Nhe1 restriction enzymes to form CYP24A1-FLASH. The promoter sequence was confirmed by Sanger Sequencing and by aligning it to the reference sequence from the Eukaryotic Promoter Database. The TK-Renilla plasmid purchased from Promega serves as an internal control for all dual-luciferase reporter assays. Under the HSV-thymidine kinase promoter, the expression of Renilla luciferase is constitutively active to normalize all firefly luciferase values. The VDR expression plasmid was generated by cloning the mature VDR transcript using first-strand cDNA synthesized using RNA isolated from HNECs. The VDR transcript was then inserted into the parental vector, pci-neo (Promega plasmid #E1841) using XhoI and SalI restriction enzymes. The VDRv1 expression vector was transiently transfected into HEK293 cells and subsequent western blotting revealed a band slightly above the 55kDa marker, representing VDR. The VDR K>R and K>Q mutants were generated using the QuikChange II site-directed mutagenesis kit by Agilent Technologies and all sequences were confirmed containing the appropriate changes.

### Dual-luciferase reporter assay

All luciferase assay data was collected using the Promega dual-luciferase reporter assay system. 24hrs post-seeding, HEK293 cells were transiently transfected with CYP24A1-FLASH (300ng), TK-Renilla (100ng) and either the empty vector control, pci-neo (160ng) or the VDR expression vector (200ng) in 1:1 molar ratio for 6hrs. 48hrs post-transfection, following any drug treatments, cells were harvested using Passive Lysis Buffer. Luciferase Assay Reagent II (LAR II) was added to the cell lysates to generate the firefly luminescent signal. The Stop & Glo Reagent quenches the firefly luminescence and generates the Renilla luminescent signal. Both signals are measured over a five-minute period and the highest peak is recorded using the Vitl Lu-mini Single Tube Luminometer. For each sample, the ratio of firefly signal over Renilla signal is calculated and compared to the negative control, which is arbitrarily set to one. All values are displayed in terms of fold-changes from the control sample.

### Mammalian Two-Hybrid System

The Promega CheckMate Mammalian Two-Hybrid System was utilized to measure the interaction between VDR and RXR. The system contains three expression vectors: PG5luc, pACT, and pBIND. PG5luc contains GAL4 binding sites upstream of the firefly luciferase gene. The pACT vector contains the herpes simplex virus VP16 activation domain upstream of a multiple coning region. The pBIND vector contains the GAL4 DNA-binding domain upstream of a multiple cloning region as well as Renilla luciferase under the control of the SV40 promoter, to serve as an internal control. The open reading frame of VDR and RXR were originally cloned from HNEC cDNA into pci-neo expression vectors. Then, both VDR and RXR were subcloned from their parental vectors (pci-neo), using primers specific to VDR or RXR. VDR, RXR, and the Two-Hybrid plasmids, pACT and pBIND, were digested with restriction enzymes Xba1 and Not1. VDR was then ligated to pBIND to create VDR-pBIND and RXR was ligated to pACT to create RXR-pACT. 24hrs post-seeding, HEK293 cells were transiently transfected with PG5luc (300ng), and 1:1 molar ratio of VDR-pBIND (100ng) and RXR-pACT (90ng) for 6hrs. 48hrs post-transfection, following appropriate drug treatments, cells were lysed and firefly and Renilla luciferase activity was measured using the Promega dual-luciferase reporter assay system kit.

### Generation of a lentiviral cell line for overexpression of His-tagged VDR

Vectors used for lentiviruses were pLVX-TRE3G and pLVX-EF1a-tet3G (Takara). VDR was sub-cloned from pci-neo vector into the pLVX-TRE3G lentiviral vector multiple cloning site and a 6-histidine tag was placed before the stop codon to create the vector pLVX-TRE3G-VDRv1-His. Lentiviruses were produced using Lenti-X Packaging Single Shots (Takara), both pLVX-EF1a-tet3G and pLVX-TRE3G-VDRv1 viruses were produced. Lenti-X 293T cells (Takara) were transfected and media was collected 48hrs after transfection.

Viral supernatants were concentrated using the Lenti-X concentrator following manufacturers protocol (Clontech). Derivation of pLVX-HEK293-VDRv1-His: HEK293 (ATCC) cells were infected with both concentrated transactivator (pLVX-EF1a-tet3G) and pLVX-TRE3G-VDRv1 viruses using polybrene (4ug/mL). The following day, cells were split into multiple plates and double selected with puromycin (1ug/mL) and hygromycin (200ug/mL). Puro/hygro resistant clones were picked and grown under continued puro/hygro selection. Clones were tested for doxycycline-induction of VDRv1-His, and for doxycycline/aD3 induced activation of our CYP-FLASH reporter. Clones were labelled, banked, and used in future experiments.

### Silver staining

pLV-HEK293-VDR-His cells were induced with doxycycline at 0.1ug/mL for 24hrs to induce expression of His-tagged VDR. Cells were pre-treated with 1uM SAHA followed by treatment with 100nM aD3 or DMSO for control for 24hrs. Cells were chemically crosslinked using 1% formaldehyde for 15mins followed by quenching with 0.125 M glycine. Cells were then lysed in denaturing conditions using guanidine buffer. The His-tag pull down was performed using Thermo Fisher Scientific HisPur Cobalt Resin for 1hr. Following the His-tag pull down, the crosslinks were reversed by heating samples at 98°C for 10mins. Samples were resolved on a Thermo Fisher Scientific Novex Tris-Glycine Protein Gel, 8-16%. Total protein was stained using Thermo Fisher Scientific Pierce Silver Stain to visualize bands.

### Coomassie staining for mass spectrometry

pLV-HEK293-VDR-His cells were induced with doxycycline at 0.1ug/mL for 24hrs to induce expression of His-tagged VDR. Cells were pre-treated with 1uM SAHA followed by treatment with 100nM aD3 or DMSO for control for 24hrs. Cells were chemically crosslinked using 1% formaldehyde for 15mins followed by quenching with 0.125 M glycine. Cells were then lysed in denaturing conditions using guanidine buffer. The His-tag pull down was performed using Thermo Fisher Scientific HisPur Cobalt Resin for 1hr. Following the His-tag pull down, the crosslinks were reversed by heating samples at 98°C for 10mins. Samples were loaded onto a Novex Tris-Glycine Mini Protein gel (10% / 1mm thick) and resolved ∼2cm into the gel. The gel was then stained with Coomassie Blue (Thermo Fisher Scientific) for 30 mins followed by 1hr of de-staining in 10% acetic acid. The entire gel was packaged and sent to the Wistar Proteomics Institute for mass spectrometry analysis.

## RESULTS

### VDR activates the VDR reporter upon treatment with aD3

The CYP24A1 promoter is a useful reporter for VDR transcriptional activity, as it is known to be regulated by VDR and contains five VDRE sites (Jones et al., 2012). An existing luciferase reporter vector was modified (TOP-FLASH) and the CYP24A1 promoter was cloned upstream of the firefly luciferase coding sequence, to create the VDR reporter vector, CYP24A1-FLASH. Using a dual-luciferase reporter assay to measure VDR transcriptional activity, VDR activity significantly increases in response to aD3 in HEK293 cells (Fig. 1). Briefly, cells were transfected with CYP24A1-FLASH, TK-Renilla internal control reporter, and either empty vector (pci-neo) or the VDR expression plasmid. Following transfection, cells were treated with either vehicle (DMSO) or 100nM aD3. In the absence of aD3, there is no activation of the VDR reporter vector. Upon treatment with aD3, there is a slight increase in VDR activity in cells transfected with empty vector (5.89-fold), representing endogenous VDR activity. In cells transfected with VDR, there is a significant increase in VDR activity upon treatment with aD3 compared to the control (109.43-fold, p=<0.0001). This demonstrates that this VDR reporter vector is a useful tool to analyze VDR transcriptional activity.

**Figure 1:**
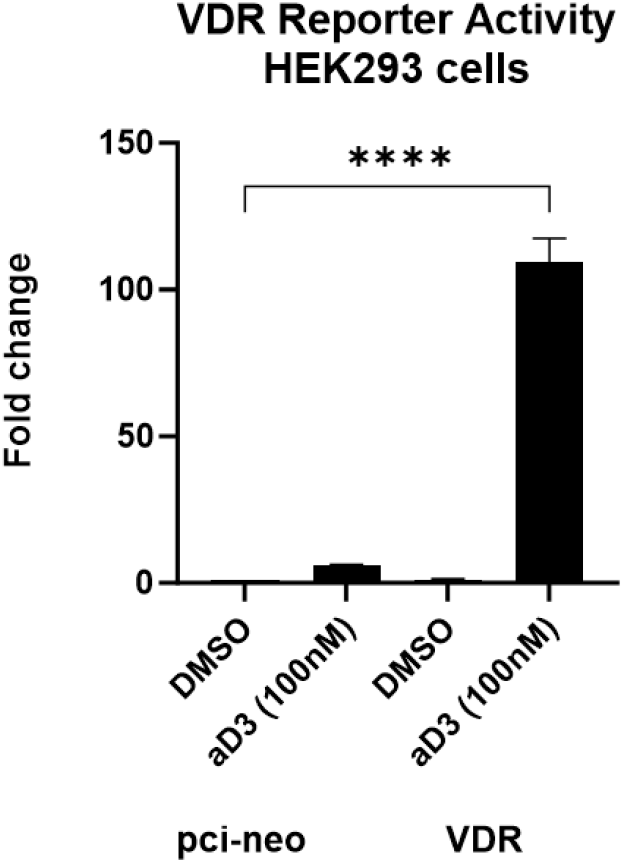
VDR reporter plasmid. HEK293 cells were transfected with CYP24A1-FLASH, control reporter TK-Renilla (100ng), and empty vector, pci-neo or VDR. Cells were then treated with vehicle, DMSO, or aD3 (100nM) for 24hrs before dual luciferase values were collected. There is a slight increase in VDR activity in the pci-neo transfected cells upon treatment with aD3, representing the endogenous VDR in the cells. There is a significant increase in VDR activity in VDR transfected cells upon treatment with aD3 (t(8) = 13.62, p=<0.0001****).

### Chemical inhibition of class I, II, and IV lysine deacetylases results in decreased VDR transcriptional activity of exogenous VDR but increased activity of endogenous VDR

To investigate whether VDR transcription is affected by lysine acetyltransferase activity, dual-luciferase reporter assays were performed to measure VDR transcriptional activity in a hyperacetylated cellular environment achieved via chemical inhibition of KDACs. The drug, SAHA (suberoylanilide hydroxamic acid), was utilized to inhibit KDACs, which catalyze the removal of lysine acetyl modifications. SAHA inhibits all eleven class I, II, and IV KDACs by binding to their active site, but does not inhibit class III KDACs (sirtuins). HEK293 cells were transfected with CYP24A1-FLASH, TK-Renilla control reporter, and either empty vector (pci-neo) or the VDR expression vector. Following transfection, cells were first pre-treated with SAHA to inhibit KDACs and allow for the accumulation of any lysine acetyl modifications followed by aD3 treatment (100nM) to activate VDR. In cells expressing exogenous VDR, there is a significant decrease in VDR activity when treating with aD3 and increasing amounts of SAHA compared to cells treated with aD3 alone (Fig. 2) (p=<0.0001). Treatment with aD3 results in an ∼89-fold increase compared to the control, whereas treatment with aD3 and SAHA (5uM) results in only a ∼13-fold increase compared to the control. Intriguingly, the opposite effect is observed in cells transfected with the empty vector control. Upon treatment with aD3 and increasing amounts of SAHA, there is a significant increase in VDR activity compared to treatment with aD3 alone. This suggests that the endogenous VDR behaves differently than the exogenously expressed VDR.

**Figure 2:**
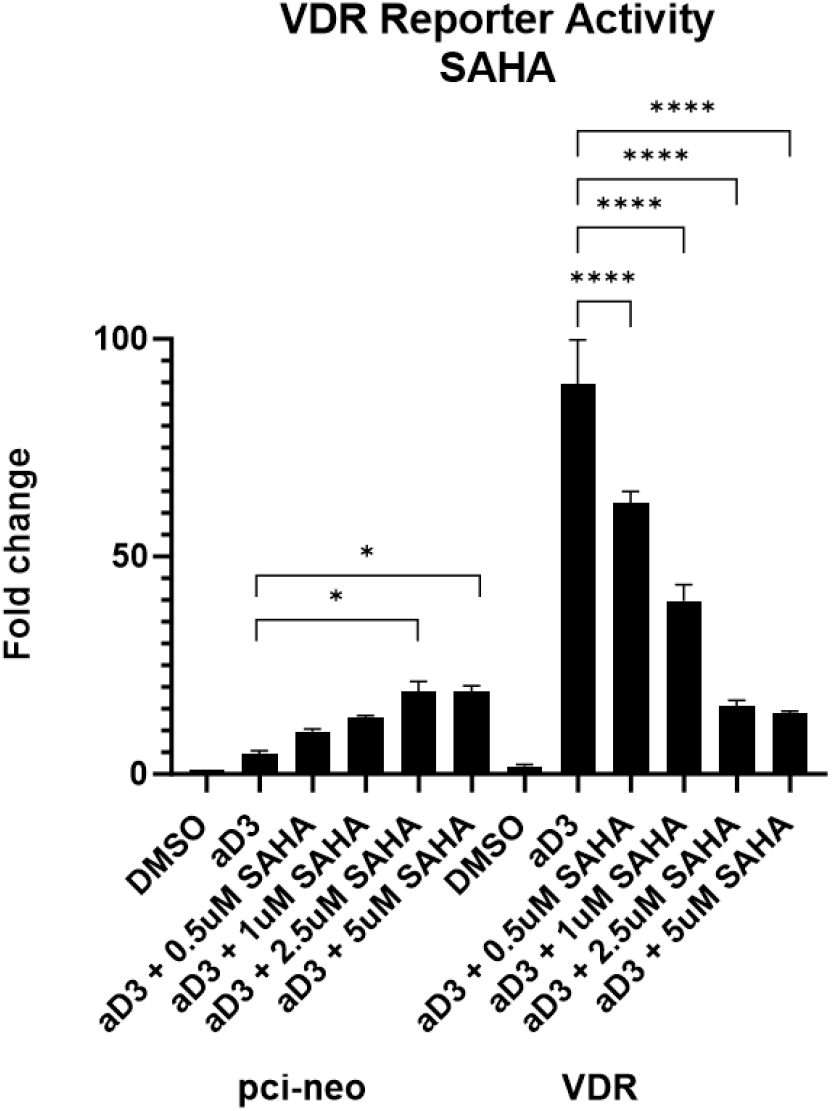
Inhibition of KDAC activity. HEK293 cells were transfected with CYP24A1-FLASH, control reporter TK-Renilla, and empty vector, pci-neo or VDR. Cells were then treated with SAHA (0.5, 1, 2.5, 5uM) for 18hrs to create a hyperacetylated environment. Cells were then treated with aD3 (100nM) for 24hrs. There is a significant increase in VDR activity in pci-neo transfected cells when treated with 2.5 and 5uM of SAHA + aD3 compared to treatment with aD3 alone (p= 0.469*, 0.0459*, respectively). There is a significant decrease in VDR activity in VDR transfected cells when treated with increasing amounts of SAHA (p=<0.0001****).

### Endogenous VDR in the HEK293 cell line carries a lysine mutation

To address why the endogenous VDR in the HEK293 cells responds differently to KDAC inhibition compared to exogenous VDR, we decided to sequence the endogenous VDR transcript in the HEK293 cell line. Briefly, RNA was isolated, followed by oligo d(t)-primed cDNA synthesis and amplification using VDR-specific primers. The endogenous VDR PCR-product was ligated into the pci-neo parental vector and transformed into competent cells. Several colonies were expanded, and clones were sent out for sanger sequencing by a third party (Azenta). After comparing each sequence to the reference wildtype VDR sequence, five out of the seven clones contain a DNA substitution of an Adenine (A) to a Guanine (G). This substitution results in an amino acid change at residue 45 from lysine to an arginine (Fig. 3). Lysine 45 is in the DNA binding domain (DBD) of VDR and directly interacts with DNA. Unlike lysine, arginine residues are not capable of undergoing acetylation, meaning residue 45 in the endogenous VDR cannot be acetylated, whereas the exogenously expressed VDR can.

**Figure 3:**
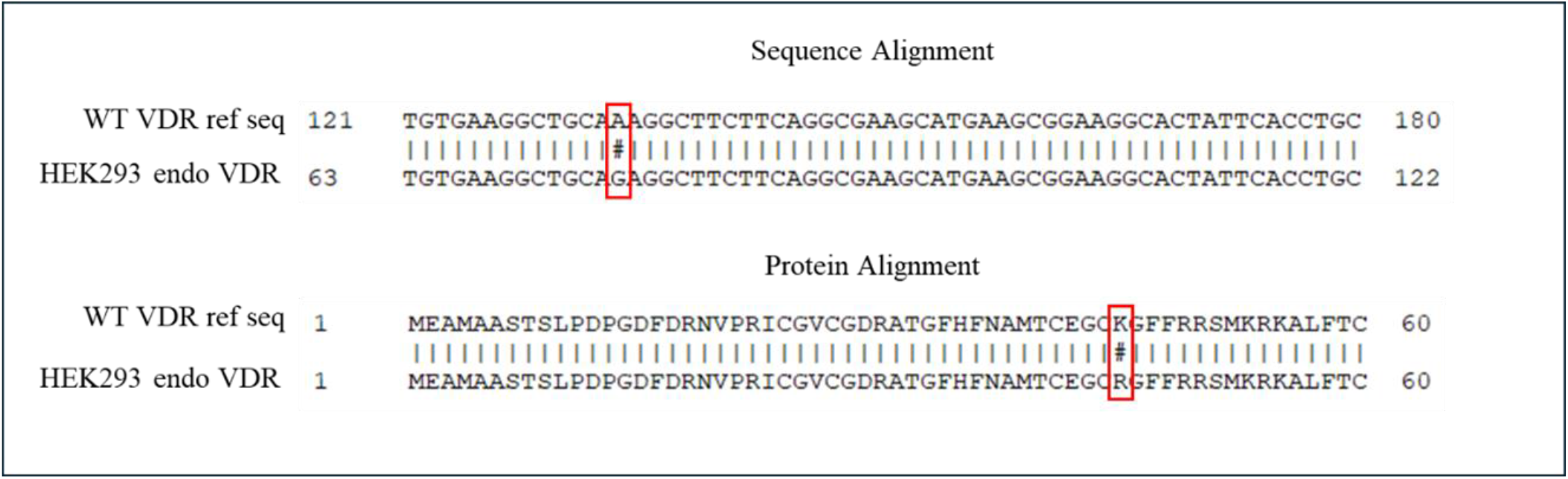
Endogenous VDR in the HEK293 cells contain a lysine to arginine mutation. Total RNA was isolated from the cells and converted into cDNA. Endogenous VDR was amplified via PCR using VDR-specific primers. VDR was then cloned into a parental pci-neo vector and transformed into competent cells. Following further expansion of colonies, seven clones were sent for Sanger Sequencing and sequences were compared back to the wildtype reference VDR sequence. Five out of seven clones contain a DNA substation of adenine (A) to guanine (G), resulting in a change from lysine (K) to arginine (R) at residue 45.

### Acetylation site predictor and site-directed mutagenesis

To investigate the consequence of lysine mutations we performed site-directed mutagenesis to create several lysine mutants of the VDR. We utilized the acetylation set enrichment-based software (ASEB), that searches for commonly acetylated motifs across a protein of interest to identify potentially modifiable residues. After running the program analysis on the VDR protein sequence, three lysine residues across VDR were predicted to undergo acetylation: K45, and K109. We also utilized a second acetylation site predictor, GPS-PAIL by the Cuckoo Workgroup. This software also predicted that lysine 109 may be a target of acetylation. For each residue, we created two mutants. A lysine (K) to arginine (R) mutant (K>R) and a lysine (K) to glutamine (Q) mutant (K>Q). The K>R substitution stabilizes the positive charge at the residue’s location, conserving the local electrostatic interactions and protein function, while abolishing its capacity for acetylation and other potential post-translational modifications. For our studies, we used the K>R mutants to represent pseudo-deacetylated lysine. The K>Q mutants represent pseudo-acetylated lysine, as the positive charge is no longer present. Glutamine’s functional group contains a nitrogen and a double-bonded oxygen, similar to the nitrogen and double-bonded oxygen of an acetyl-lysine. In addition, glutamine maintains a neutral charge, like an acetylated lysine. All mutant expression vectors were sequenced and confirmed to contain the desired mutation.

### VDR K45R has diminished aD3-mediated activation whereas K45Q is unresponsive to aD3

Compared to WT VDR, the deacetylated mimic, K45R, is significantly less active in response to aD3 (∼54-fold increase in WT VDR compared to ∼15-fold increase in VDR K45R). However, unlike WT VDR, creating a hyperacetylated environment via SAHA results in an increase in VDR activity in the K45R mutant (∼19-fold increase in WT VDR compared to ∼23-fold increase in VDR K45R), consistent with our findings from HEK293 cell endogenous VDR that also carries the K45R mutation (Fig. 4). Furthermore, the acetylated mimic, K45Q, is unresponsive to aD3-mediated activation. Treatment with aD3 alone or aD3 in combination with SAHA does not activate VDR K45Q, resulting in only a ∼1-fold increase in both conditions compared to the control.

**Figure 4:**
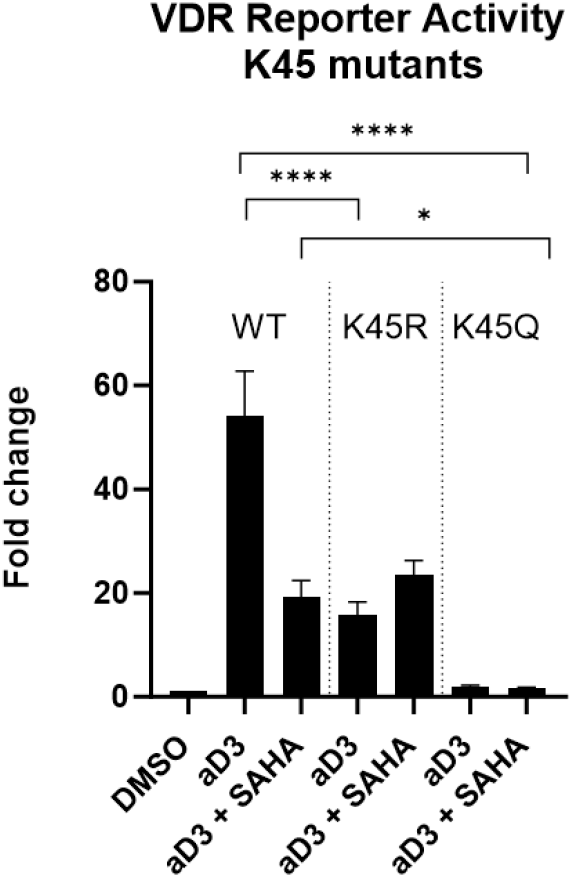
VDR K45 mutant assay. HEK293 cells were transfected with the VDR reporter, CYP24A1-FLASH, (300ng), control reporter TK-Renilla (100ng), and VDR (WT, K45R, K45Q – 200ng) for 6hrs. Cells were then treated with SAHA (1uM) for 18hrs to create a hyperacetylated environment. Cells were then treated with aD3 (100nM) for 24hrs. There is a significant decrease in VDR activity in response to both aD3 and aD3 + SAHA in the mutants compared to WT VDR (F=33.13, p=<0.0001****).

### VDR K109R has diminished aD3-mediated activation that can be restored with K109Q

To investigate the ramifications of pseudo-acetylation and deacetylation of lysine 109 on VDR activity, we performed dual-luciferase reporter assays using the K109R and K109Q mutants to measure their activity under baseline conditions as well as in a hyperacetylated environment. Compared to wildtype VDR, the de-acetylated mimic, K109R, is less active in response to treatment with aD3 (∼95-fold increase in WT VDR compared to a ∼32-fold increase in VDR K109) (Fig. 5). By creating a hyperacetylated environment using SAHA, VDR K109R, whose residue’s positive charge is stabilized and cannot undergo acetylation, loses activity compared to the wildtype VDR (∼40-fold increase in WT VDR compared to ∼26-fold increase in VDR K109R). Alternatively, mimicking an acetylated residue by neutralizing the positive charge in the VDR K109Q mutant, activity in response to aD3 is restored back to the level of wildtype VDR and there is no significant difference between WT VDR and VDR K109Q in response to aD3 treatment. VDR K109Q still exhibits a reduction in its activity in response to KDAC inhibition.

**Figure 5:**
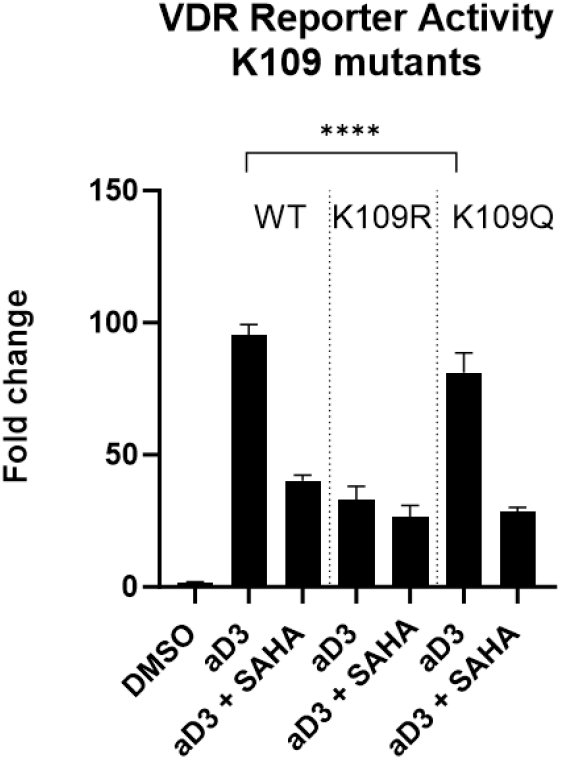
VDR K109 mutant assay. HEK293 cells were transfected with the VDR reporter, CYP24A1-FLASH, (300ng), control reporter TK-Renilla (100ng), and VDR (WT, K109R, K109Q – 200ng) for 6hrs. Cells were then treated with SAHA (1uM) for 18hrs to create a hyperacetylated environment. Cells were then treated with aD3 (100nM) for 24hrs. There is no significant decrease in VDR K109Q activity in response to aD3 compared to WT VDR activity in response to aD3. There is a significant decrease in VDR K109R activity in response aD3 compared to WT VDR activity in response to aD3 (p=<0.0001 ****).

### Inhibition of lysine methyltransferases decreases VDR transcriptional activity

Given that post-translational lysine methylation often occurs at the same lysine residues that are acetylated, typically yielding opposite effects on protein activity, we decided to investigate if VDR activity can be modulated through inhibition of methyltransferases. To do so, we used chemical inhibitors of several lysine specific methyltransferases (KMTs) to prevent the addition of methyl groups onto lysine residues.

KMTs catalyze the methylation of lysine residues using the co-factor S-adenosyl-l-methionine (SAM) as the methyl donor. We selected three drugs to inhibit three different families of KMTs. UNC0642 inhibits KMT G9a/GLP, which has been shown to mono- and di-methylate various transcription factors (Huang et al., 2010; Ling et al., 2012; Pess et al., 2008). UNC0379 inhibits SETD8 KMT which is responsible for mono-methylating various non-histone proteins (Shi et al., 2007; Takawa et al., 2012). GSK126 inhibits EZH2, which has been shown to methylate various transcription factors, including a nuclear receptor (Lee et al., 2012). For this set of experiments, following transfection of the VDR reporter, TK-Renilla control reporter, and VDR, cells were pre-treated with the KMT inhibitors to prevent the addition of methyl groups onto lysine residues, followed by treatment with aD3 to activate VDR. Upon inhibition of each individual KMT and treatment with aD3, there is a significant decrease in VDR transcriptional activity compared to treatment with aD3 alone (Fig. 6). Suggesting that transferring of methyl groups to lysine residues is necessary to reach maximal aD3-mediated VDR transcriptional activity.

**Figure 6:**
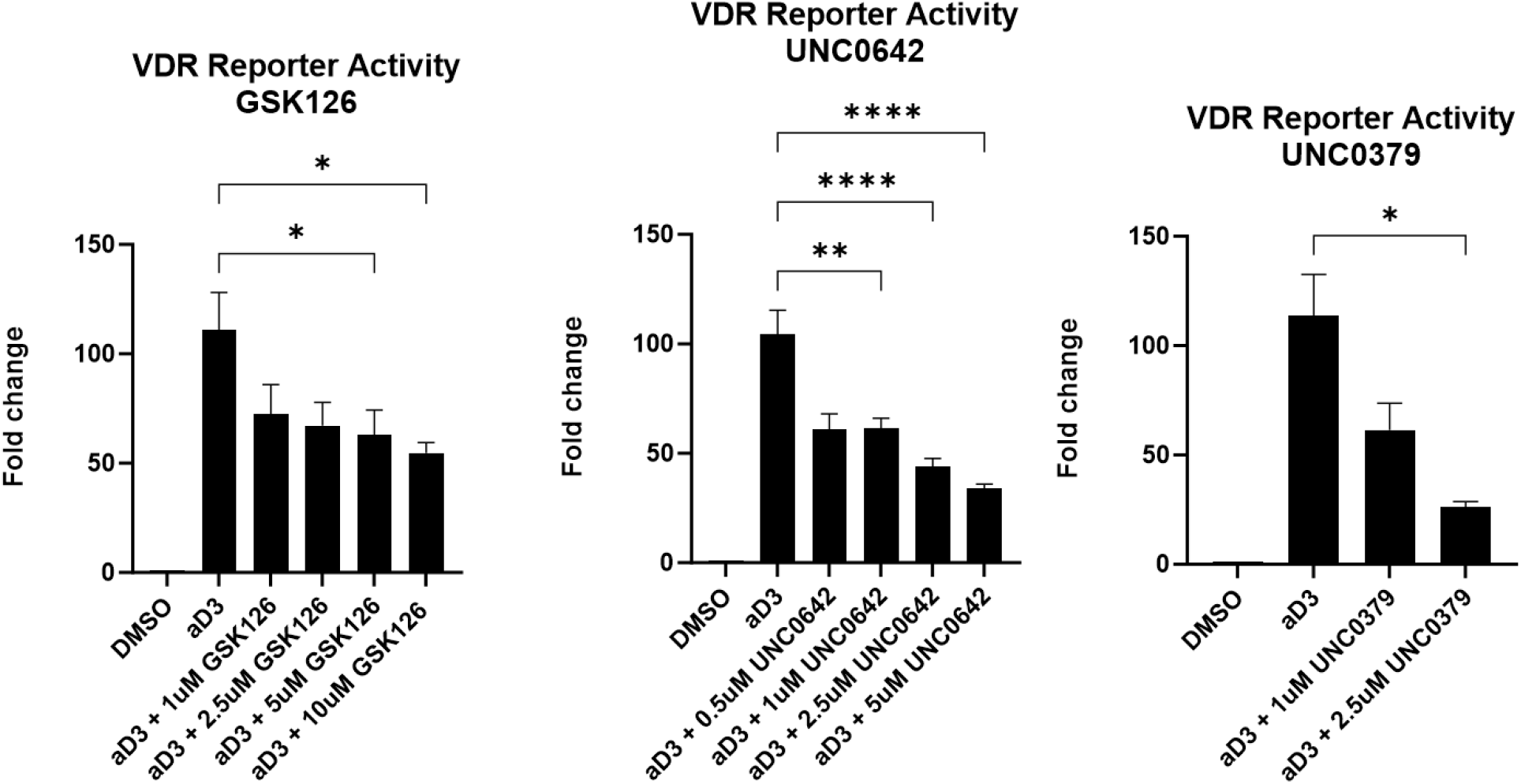
KMT inhibition. HEK293 cells were transfected with the VDR reporter, CYP24A1-FLASH, (300ng), control reporter TK-Renilla (100ng), and VDR (200ng) for 6hrs. Cells were then treated with either GSK126 (1, 2.5, 5, 10uM), UNC0642 (0.5, 1, 2.5, 5uM) or UNC0379 (1, 2.5uM) for 18hrs to inhibit methyltransferases from methylating VDR. Cells were then treated with aD3 (100nM) for 24hrs. There is a significant decrease in VDR activity in response to the KMT inhibitor drugs with aD3 compared to treatment with aD3 alone. (GSK126: F=10.24, p=0.0005***, UN0642: F=30.21, p=<0.0001****, UNC0379: F=19.29, p=0.007**).

### The effect of chemical inhibition of lysine methyltransferases and lysine deacetylases on VDR’s affinity for RXR

Given that we can modulate VDR transcriptional activity by preventing the addition of methyl marks or by stabilizing acetyl marks, we wanted to investigate if this change in activity is a result of altering the ability of VDR to bind to its heterodimeric partner, RXR. To do so, a mammalian Two-Hybrid system was used. This system utilizes a DNA binding domain and a transcriptional activation domain, which when brought together form a functional transcriptional activator. We cloned VDR to the DNA binding domain and RXR to the activation domain. When VDR and RXR interact, the DNA binding domain and activation domain will recruit the required protein machinery to transcribe the firefly luciferase reporter gene. Upon treatment 100nM aD3, there is no significant change in VDR:RXR interaction, which is to be expected as the two are bound in both the presence and absence of aD3 (Fig. 7). When treating with methyltransferase inhibitor, GSK126, where VDR is less transcriptionally active, there is no significant change in VDR:RXR interaction. Upon treatment with KDAC inhibitor, SAHA, where VDR is also less transcriptionally active, there is a significant increase in VDR:RXR interaction. These results suggest that the VDR:RXR interaction is stabilized while being held in a transcriptionally repressive state.

**Figure 7:**
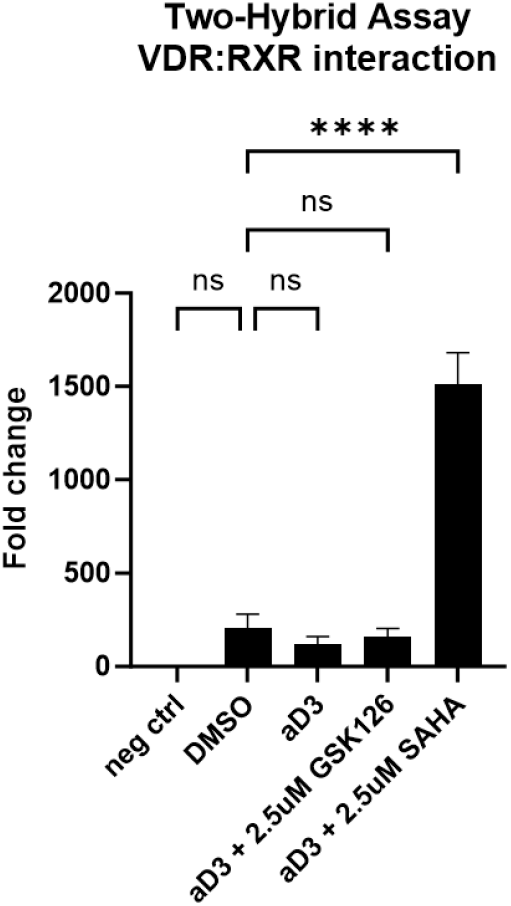
Two-hybrid assay. HEK293 cells were co-transfected with PG5-luc, VDR-pBIND, and RXR-pACT. Cells were pre-treated with either GSK126 (2.5uM) or SAHA (2.5uM) for 18hrs. Cells were then treated with aD3 (100nM) for 24hrs. There is no significant change in VDR:RXR interaction upon addition of aD3 or aD3 + GSK126. There is a significant increase in VDR:RXR interaction upon treatment with aD3 + SAHA (p=<0.0001 ****).

### Chemical crosslinking and denaturing VDR pulldowns reveal differential banding patterns in cells treated with a deacetylase inhibitor

Given that the interaction of VDR and RXR is stabilized in a hyperacetylated cellular environment, even though VDR activity is repressed, VDR activity could be modulated by a failure of co-repressors to dissociate or preventing the association of co-activators. To investigate this possibility, we performed chemical crosslinking, denaturing pulldowns, and total protein staining to see if we could detect differences in banding patterns across conditions. To do so, a doxycycline-inducible lentiviral HEK293 cell line (pLV-HEK293-VDR-His) was engineered so that upon treatment with doxycycline, a Histidine-tagged VDR is expressed. Following overexpression of His-tagged VDR, pLV-HEK293-VDR-His cells were treated with either DMSO, aD3, or both SAHA and aD3. Before harvesting, cells were chemically crosslinked with formaldehyde to capture any proteins that were complexed with VDR. A His-tag pull down via cobalt metal affinity resin was performed in denaturing conditions to reduce nonspecific interactions. Prior to loading on a gel, the crosslinks were reversed, and silver staining was performed to stain total protein in each well (Fig. 8). We observed shifting banding patterns both with the addition of aD3 and with SAHA pre-treatment. Each of our three conditions displayed unique banding patterns, supporting our hypothesis that SAHA pre-treatment is altering the composition of VDR protein-protein complexes.

**Figure 8:**
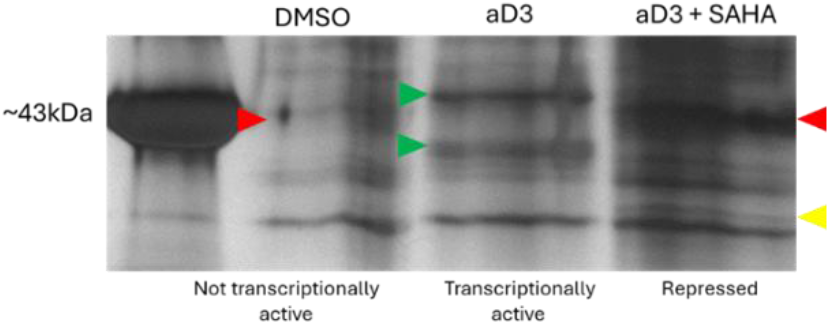
SAHA alters VDR protein-protein interactions. 24hrs after seeding, pLV-HEK293-VDR-His cells were induced with 0.1ug/mL of doxycycline to overexpress His-tagged VDR. Cells were pre-treated with SAHA (1uM) followed by aD3 (100nM). 24hrs later, cells were chemically crosslinked with formaldehyde. Cells were then harvested in denaturing conditions and a cobalt resin was used to immunoprecipitate his-tagged VDR and associated proteins. Following reversal of the crosslink and separation on a Tris-Glycine gel, total protein was stained using Thermo Scientific Silver Staining Kit. There are two bands present in the aD3 lane that are not present in the other two (green arrows). There is one band present in both the control lane and SAHA lane that is not present in the aD3 lane (red arrows). There is a band present in the SAHA lane that is not present in the other two lanes (yellow arrow).

### Mass spectrometry for protein identification and abundance in VDR crosslinked samples reveals altered protein-protein interactions

Given that we observed differential silver-stained banding patterns in VDR pull downs in HEK293 cells treated with DMSO, aD3, or aD3 with SAHA pre-treatment, we decided to analyze all three pull downs using mass spectrometry to identify differences in protein targets and abundance. Briefly, the lentiviral HEK293 cells were induced with doxycycline (0.1ug/mL) to overexpress His-tagged VDR, followed by treatment with either DMSO, aD3 (100nM), or SAHA (1uM) and aD3. Lysates were chemically crosslinked using formaldehyde followed by His-tag pull downs with cobalt metal affinity resin in denaturing conditions.

Crosslinks were reversed prior to SDS-PAGE, samples were resolved ∼0.5cm to concentrate all proteins into a single band, the gel was stained with Coomassie Blue and sent to Wistar Institute for mass spectrometry and protein identification. Mass spectrometry identified the proteins present in each condition as well as their relative abundance. We plotted the normalized label-free quantification (lfq) intensity values to compare relative abundance of proteins between the three conditions.

We observed from the dual-luciferase reporter assays that SAHA pre-treatment inhibits aD3-mediated VDR activation. We also discovered that this repression is not resulting from the inability of VDR to bind to its heterodimeric partner, RXR, as shown by the Two-Hybrid data, where VDR and RXR binding is stabilized (Fig. 7). To further confirm these findings, we looked at the lfq intensity values from the mass spectrometry analysis, which measures normalized protein abundance across samples. We found more RXR interacting with VDR in cells pre-treated with SAHA compared to cells treated with vehicle or aD3 alone, confirming the results found from our Two-Hybrid assays (Fig. 9).

**Figure 9:**
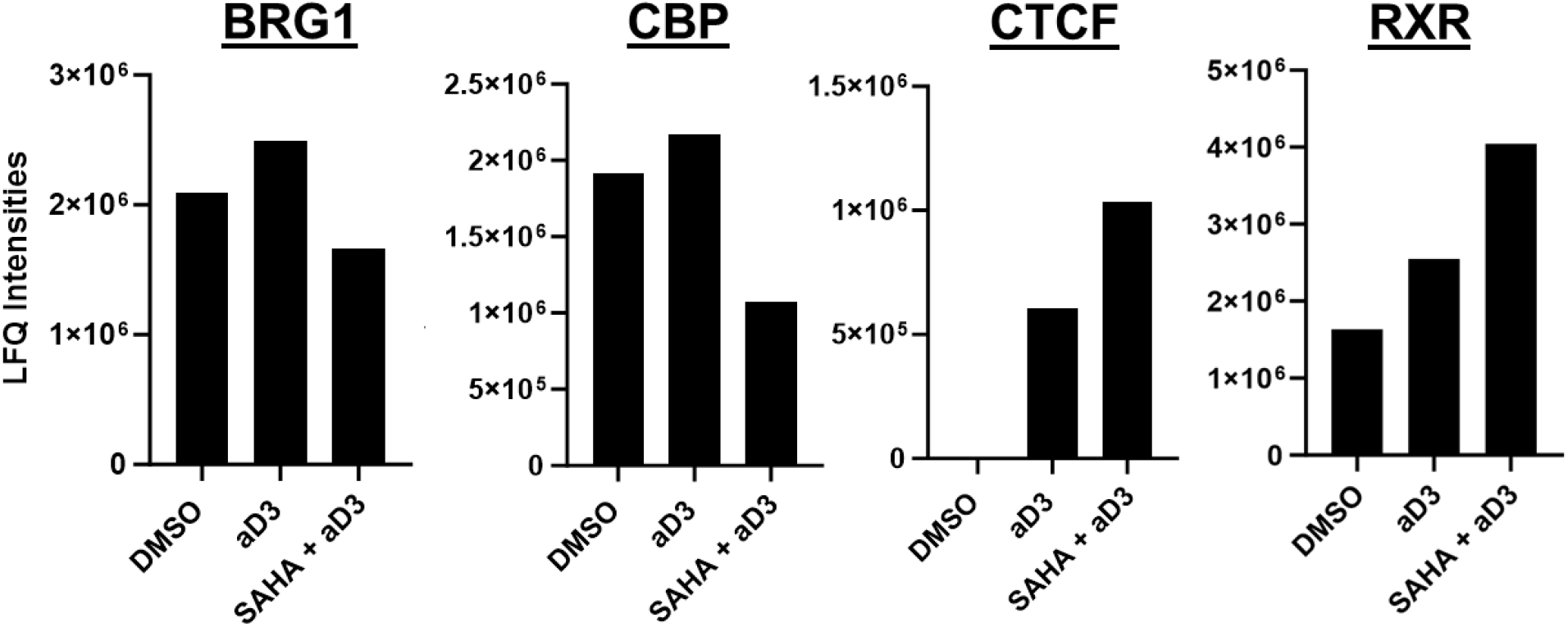
Mass spec analysis. pLV-HEK293-VDR-His cells were induced with doxycycline (0.1ug/mL) to overexpress VDR-His, followed by treatment with either control (DMSO), aD3 (100nM), or aD3 + SAHA (1uM). Cells were chemically crosslinked prior to the His-tag pull downs and resolved ∼0.5cm on SDS-PAGE to concentrate proteins from each sample into a single band. The gel was analyzed with mass spectrometry, and lfq values measuring normalized protein abundance was generated. Normalized lfq intensity values were plotted to compare relative protein abundance in each condition. The abundance of three co-activators decreases in cells treated with SAHA, where VDR is inactive: BRG1, CBP, and NCoA-5. The abundance of two co-repressors increases in cells treated with SAHA: Sin3A and CTCF.

Several co-activators are more abundant in their interaction with VDR in aD3-treated cells, where VDR is transcriptionally active, compared to the SAHA and aD3-treated cells, where VDR transcription is inhibited. For example, the BRG1 transcriptional activator is the most abundant in cells treated with aD3 alone and is least abundant in cells treated with aD3 and SAHA. Another example is the CREB binding protein, which is also most abundant in cells treated with aD3 alone and least abundant in cells treated with aD3 and SAHA. Alternatively, there are some co-repressors that are enriched in SAHA-treated cells. For example, the CCCCTC-Binding Factor (CTCF) repressor is most abundant in cells treated with both SAHA and aD3 (Fig. 9). The altered affinity for VDR exhibited by these co-activators and co-repressors after SAHA treatment demonstrate that inhibition of KDAC activity inhibits the proper formation of the VDR transcriptional complex.

## Discussion

### VDR transcriptional activity can be modulated by creating a hyperacetylated environment through inhibition of KDACs classes I, II, and IV

When treating cells with SAHA to inhibit class I, II and IV KDACs, thus creating a hyperacetylated environment, aD3-mediated VDR activity significantly decreases compared to when treating with aD3 alone. This suggests that stabilizing lysine acetyl marks has an inhibitory effect on VDR transcriptional activity. When performing the same experiment with 3TYP, which inhibits class III KDACs, the Sirtuins, there is no significant change in VDR activity (data not shown). This suggests that if VDR or the VDR transcription complex undergoes lysine acetylation, it is class I, II, and IV KDACs that may be responsible for removing the acetyl marks, not Sirtuins.

Taken together with the SAHA data, potential acetylation of VDR or the VDR complex is required for normal VDR transcriptional activity, and subsequent removal of these lysine acetyl marks may be catalyzed by class I, II, and IV KDACs.

### Site directed mutagenesis points to several residues in VDR that may undergo acetylation and/or other PTMs

The data from our dual-luciferase reporter assays demonstrate that HEK293 endogenous VDR and wild-type exogenous VDR respond oppositely to a hyperacetylated environment. Endogenous VDR activity increases when acetyl marks are stabilized, whereas exogenous VDR activity decreases. We know from sequencing endogenous VDR in the HEK293 cell-line that VDR carries a lysine to arginine (K>R) mutation at residue 45. This means that residue 45 in the endogenous VDR cannot undergo post-translational acetylation or other PTMs. This suggests that if acetylation does indeed occur on VDR, specifically at lysine 45, it may be inhibitory to VDR activity because when it is mutated to an arginine and cannot be acetylated, we no longer observe this inhibitory effect. In fact, VDR activity increases in response to KDAC inhibition in the K45R endogenous VDR, suggesting that there may be other acetylation sites on VDR or an associating protein that act positively on VDR transcriptional output.

### Potential acetylation at residue 109 may be a positive regulator of VDR activity

In the deacetylated mimic, K109R, there is a significant decrease in VDR transcriptional activity compared to wildtype (WT) VDR in response to both aD3 and aD3 plus SAHA. On the other hand, the pseudo-acetylated mimic (lysine to glutamine, K>Q), K109Q, displays activity similar to WT VDR. This suggests that potential acetylation at this residue positively affects VDR activity, because when we prevent acetylation at this site via K>R mutagenesis, VDR activity decreases. These data support the hypothesis that potential acetylation at residue 109 is a positive regulator of transcriptional activity. Given the observation that VDR K109Q loses activity in response to SAHA, like WT VDR, suggests that there may still be other lysine residues across the protein that contribute to the loss of VDR activity when acetyl marks are stabilized via KDAC inhibition.

### Potential acetylation at residue 45 may act as a negative regulator of VDR activity

The deacetylated mimic, K45R, is less responsive to aD3 compared to WT VDR. However, when stabilizing acetyl marks via SAHA, there is an increase in VDR activity in the K45R mutant. This data is consistent with our data from HEK293 endogenous VDR, which we now know is also K45R. This suggests that if residue 45 is acetylated, it is inhibitory of VDR activity. When this residue is an arginine, thus incapable of undergoing acetylation, treating with SAHA may stabilize acetyl marks on other residues that act as positive regulators of VDR activity (such as residue 109). To further support this hypothesis, the pseudo-acetylated mimic, K45Q, is totally inactive. Treatment with aD3 alone or the combination of aD3 and SAHA cannot activate VDR K45Q, further suggesting that acetylation at this residue is inhibitory to transcription. However, this still does not explain why the de-acetylated mimic is unable to restore activity back to WT VDR levels. It is possible that residue 45, when a lysine, undergoes a different PTM, potentially methylation. Residue 45 is in the DBD, in a motif that is conserved across all NHRs (CEGCKG). Other studies have reported that this exact lysine in other NHRs can be methylated. For example, when Retinoic Acid receptor (RAR) is mono-methylated at this residue, it helps to coordinate co-activators on the transcription complex and orients the DBD and LBD for activation (Huq et al., 2008). It’s possible that in WT VDR, lysine 45 undergoes methylation which allows for maximum activation in response to aD3. Acetylation at this same residue acts as an “off-switch” to inactivate VDR, which is evident from our VDR K45Q mutant data. By changing this lysine to an arginine, methylation is inhibited, which could explain why aD3-mediated transcription is reduced in the VDR K45R mutant compared to WT VDR.

### Methylation of VDR or the VDR transcription complex may positively influence VDR activity

To further investigate the potential that VDR activity can be modulated by methylation, we used drugs to inhibit various lysine methyltransferases (KMTs). By inhibiting KMTs, thus preventing them from methylating their substrates, there is a significant decrease in VDR activity, suggesting that methylation of VDR or its transcriptional complex members increases activity. Tied together with the data from the acetylation mutants, we can begin to parse apart the various PTMs VDR may carry and their effect on VDR’s transcriptional output. VDR may undergo methylation, potentially at residues 45 and 53, that helps orient the DBD of VDR to allow for maximum activation in response to aD3. It’s possible that acetylation may also occur on residue 45 and may negatively influence VDR activity. Residue 109 may undergo acetylation to positively influence VDR activity. We cannot rule out the potential that these residues may undergo other PTMs that are not acetylation or methylation, and further research needs to be conducted to explore this possibility.

### A hyperacetylated cellular environment stabilizes certain VDR protein-protein interactions while preventing others

We know from our dual-luciferase reporter assay data that when treating cells with SAHA and creating a hyperacetylated environment, VDR transcriptional activity significantly drops. We also know from our two-hybrid assay data, that treating with SAHA surprisingly stabilizes VDR and RXR. This suggests that the loss in VDR activity is not due to the prevention of VDR’s ability to bind to RXR. SAHA may result in decreased VDR transcriptional activity by stabilizing interactions with co-repressors that associate with VDR in the absence of aD3 or preventing the association of co-activators in the presence of aD3. It is known that when aD3 binds with VDR there is a dissociation of co-repressors and a recruitment of co-activators, when we silver stained total protein pulled down with our his-tagged VDR under various conditions (DMSO, aD3, aD3 + SAHA), we identified shifts in banding patterns depending on the condition. We interpret the band shifts from DMSO to aD3 as the necessary alterations in protein-protein interactions to facilitate VDR transcription. We interpret bands which persist between DMSO and aD3+SAHA as potential co-repressors which were unable to dissociate from the VDR complex. For example, the band that is minimally present in the DMSO lane and greatly stabilized in the SAHA lane may be a co-repressor that is stabilized on the VDR transcription complex in the presence of SAHA. Similarly, bands which appear in aD3 but fail to appear in aD3+SAHA we interpret as co-activators that are unable to associate with the VDR complex. For example, there are two bands that are only present in the aD3 lane, where VDR is highly transcriptionally active that may be co-activators. Finally, there is a band that is present only in the SAHA lane, which could possibly be a novel binding partner of VDR. Ultimately, these data demonstrate that preventing the removal of lysine acetylation modifications will alter VDR protein-protein interactions when responding to aD3.

### Mass spectrometry for protein identification and abundance in VDR crosslinked samples in control, aD3, and SAHA conditions

Mass spectrometry for total protein identification was performed on cross-linked VDR-his pull downs in three different conditions: control (DMSO), where VDR is inactive, aD3, where VDR is highly transcriptionally active, and aD3 plus SAHA, where VDR transcription is repressed. Using the label-free quantification (lfq) intensity values, which represents normalized protein abundance, we found various proteins that were pulled-down with VDR in different amounts in each condition which may shed light on why SAHA prevents maximal activation of VDR transcription. There are three co-activators that are more abundant in the aD3-treated sample compared to the SAHA-treated sample. For example, BRG1 is a transcriptional activator that acts as a chromatin remodeling complex and facilitates transcription factor binding. BRG1 has already been shown to activate several other nuclear hormones including peroxisome proliferator activated receptor, estrogen receptor, and androgen receptor (Debril et al., 2004; Ichinose et al., 1997; Link et al., 2005). The amount of BRG1 pulled-down with VDR was most abundant in cells treated with aD3 alone and is least abundant in cells treated with aD3 and SAHA. CREB binding protein (CBP), a 40kDa co-activator, is also most abundant in VDR pulldowns performed on cells treated with aD3 alone and least abundant in cells treated with aD3 and SAHA. Previously, CBP has been shown to enhance other nuclear hormone transcriptional activity, specifically for the glucocorticoid receptor, estrogen receptor, and androgen receptor (Aarnisalo et al., 1998; Dutertre & Smith, 2003; Sheppard et al., 1998). Studies have also shown CBP potentiates VDR signaling at specific promoters together with the recruitment of another co-activator, SRC-1 (Castillo et al., 1999). Overexpression of CBP has also been shown to alleviate VDR repression by the co-repressor YY1 by preventing its binding to VDR (Raval-Pandya et al., 2001). We identified a co-repressor that was enriched in VDR pulldowns in cells treated with SAHA that may play a role in repressing VDR in this hyperacetylated state. The CCCCTC-Binding Factor (CTCF) repressor is most abundant in cells treated with both SAHA and aD3. This zinc finger protein works as a transcription factor by blocking communication between enhancers and promoters and recruits other co-repressors.

Specifically, CTCF has been shown to prevent binding of the estrogen receptor to chromatin, thus decreasing transcriptional activity (Fiorito et al., 2016). Similarly, in the thyroid receptor, CTCF recruits another co-repressor, Sin3A, and together they repress transcription (Lutz et al., 2000). Taken together, these data demonstrate that inhibition of KDAC activity alters affinity of members of the VDR transcriptional complex.

Our investigations in VDR transcription have demonstrated that KDAC and KMT activities influence aD3-mediated transcriptional activation of the VDR. Inhibition of these enzymes can positively or negatively alter the transcriptional output of the VDR. We have shown that inhibition of class I, II, and IV KDACs will impact the ability of the VDR transcriptional complex to form properly and therefore decrease overall transcriptional output. We have also demonstrated, through site-directed mutagenesis of VDR lysines, that pseudo-acetylation of certain lysines can either decrease or increase the transcriptional output of the VDR in response to aD3. Taken together these findings highlight an intriguing additional layer of regulation to VDR transcription. The simplest explanation for our observations in our SAHA and site-directed mutagenesis experiments is that the VDR is being directly acetylated at certain lysines and this alters VDR transcriptional activity, this would need to be confirmed via mass spectrometry. It is also possible that accessory proteins such as co-repressors and co-activators are directly acetylated/de-acetylated, and this is necessary for proper dissociation and recruitment of VDR complex members. Finally, we cannot rule out that the SAHA effects on VDR transcription may be due to an indirect pathway such as alteration of a separate signaling network which indirectly influences the transcription of VDR through altering stability/availability of VDR co-repressors/co-activators.

Understanding novel ways that cells regulate VDR transcriptional activity could be useful in treating various diseases. For example, being able to fine tune VDR activity with SAHA, which is already clinically approved for humans, may prove useful in certain types of cancers such as myeloid leukemia and papillary thyroid cancer where VDR is overexpressed. In addition, it may prove meaningful to know if cancer patients are harboring VDR lysine mutations to understand if a cancer has upregulated or downregulated VDR transcription. Overall, understanding the mechanism on how VDR can be regulated, whether it be through the activity of specific KATs or KDACs, or the presence or absence of certain co-activators or co-repressors, can help us understand how VDR and vitamin D signaling may be contributing to various diseases and cancers.

